# souporcell: Robust clustering of single cell RNAseq by genotype and ambient RNA inference without reference genotypes

**DOI:** 10.1101/699637

**Authors:** Haynes Heaton, Arthur M. Talman, Andrew Knights, Maria Imaz, Daniel Gaffney, Richard Durbin, Martin Hemberg, Mara Lawniczak

## Abstract

Methods to deconvolve single-cell RNA sequencing (scRNAseq) data are necessary for samples containing a natural mixture of genotypes and for scRNAseq experiments that multiplex cells from different donors^1^. Multiplexing across donors is a popular experimental design with many benefits including avoiding batch effects^2^, reducing costs, and improving doublet detection. Using variants detected in the RNAseq reads, it is possible to assign cells to the individuals from which they arose. These variants can also be used to identify and remove cross-genotype doublet cells that may have highly similar transcriptional profiles precluding detection by transcriptional profile. More subtle cross-genotype variant contamination can be used to estimate the amount of ambient RNA in the system. Ambient RNA is caused by cell lysis prior to droplet partitioning and is an important confounder of scRNAseq analysis^3^. Souporcell is a novel method to cluster cells using only the genetic variants detected within the scRNAseq reads. We show that it achieves high accuracy on genotype clustering, doublet detection, and ambient RNA estimation as demonstrated across a wide range of challenging scenarios.

---

The ability to demultiplex mixtures of genotypes from droplet-based scRNAseq protocols, e.g. drop-seq^4^ or 10x Genomics^5^, is important because mixed sample scRNAseq is a powerful experimental design that reduces costs per donor, controls for technical batch effects, and provides information on both cross-genotype doublets and the amount of ambient RNA in the experiment. Until recently, a genotype reference obtained via whole genome or exome sequencing has been required for each multiplexed individual prior to cell-sample categorization. We present souporcell, a method to cluster cells by genotype, call doublet-cell barcodes, and infer the amount of ambient RNA in the experiment without the use of a genotype reference. We compare our method to demuxlet^1^, the gold standard method that requires genotype information *a priori*, as well as two new tools that, like souporcell, do not require prior genetic information^6,7^. We show that souporcell not only outperforms these new methods, but also surpasses demuxlet on both cell assignment and doublet accuracy. Furthermore, souporcell is the only tool that explicitly models and estimates the amount of ambient RNA in the experiment, which is a major confounder of scRNAseq analysis with regard to both expression and genotype. Although a tool for ambient RNA quantification exists^3^, it requires prior knowledge in the form of one or more well expressed genes known to not be expressed in a particular cell type. Souporcell is freely available under the MIT open source license at https://github.com/wheaton5/souporcell.

To cluster cells by genotype, we first must measure the allele information for each cell. To achieve the most accurate clustering, it is imperative that the variant calls and allele counts are measured accurately. While other tools start from the STAR aligned bam^8^ that is produced as part of running cellranger^9^, we have found several artifacts of the STAR alignments (methods) that are a significant source of false positive variants and reference bias. Instead, we remap the reads with minimap2^10^ (Fig. 1a) which produces alignments more conducive to accurate variant calling. We call putative single nucleotide polymorphisms (SNPs) with freebayes^11^ (Fig. 1b). Next, we count alleles per cell with vartrix^12^ (Fig. 1c) which avoids reference bias due to ambiguous support such as alignment end effects. If a source of reliable common variants is available, this can be used instead of the freebayes candidate variants.

**Figure 1:**
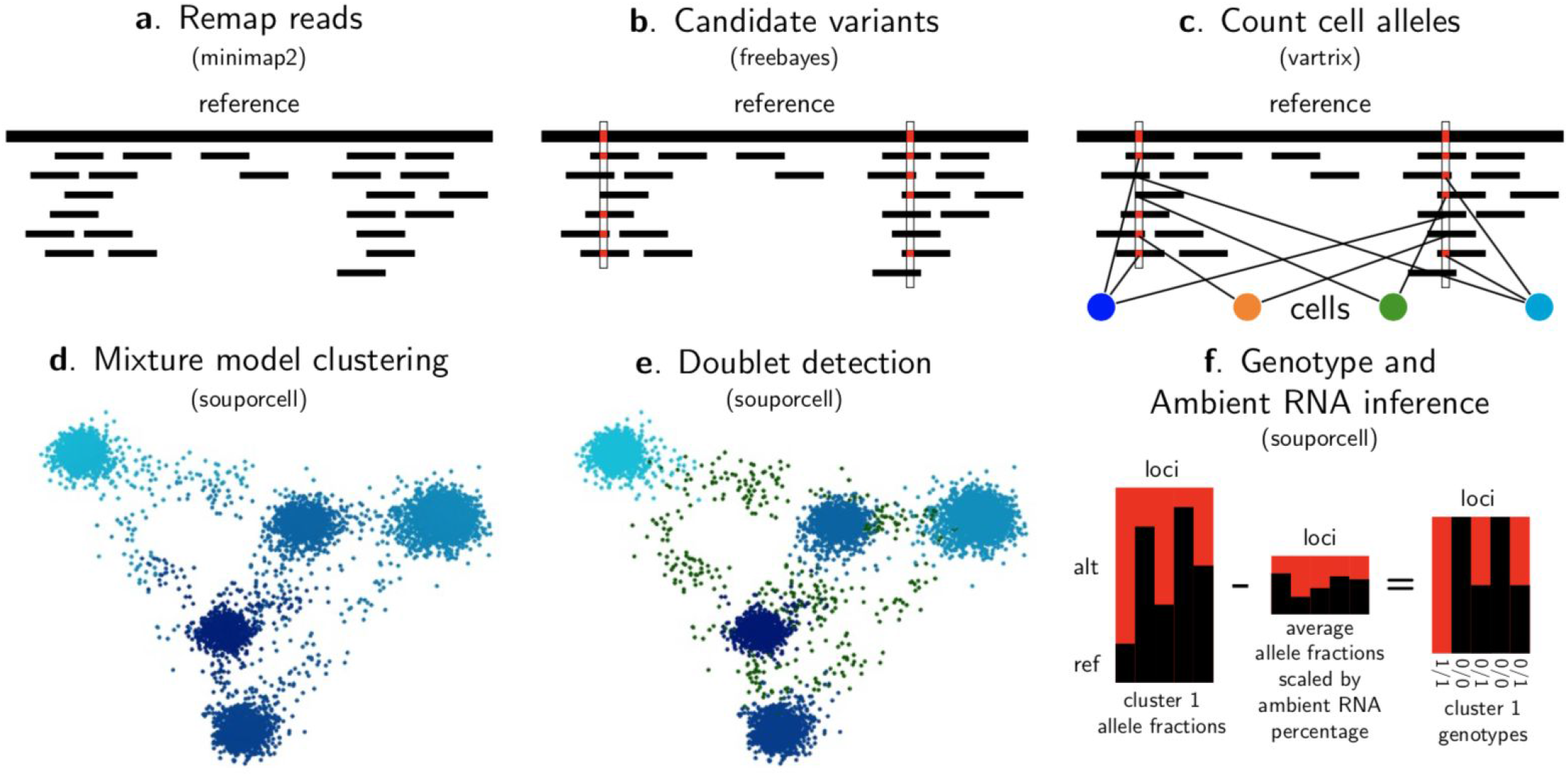
souporcell overview. **a**, We first remap the reads using minimap2 retaining the cell barcode and unique molecular identifier barcode for downstream use. **b**, We then call candidate variants using freebayes and **c**, count the allele support for each cell using vartrix. **d**, Using the cell allele support counts, we cluster the cells using sparse mixture model clustering (methods). **e**, Given the cluster allele counts, we categorize cells as doublets or singletons and excluding those doublets, **f**, we infer both the fraction of ambient RNA and the genotypes of each cluster (example for one cluster).

The clustering problem can be represented as a matrix *X* where each row represents a cell, each column represents a variant, and each element is the number of reads supporting each allele of the variant. We fit a mixture model with the cluster centers represented as the alternate allele fraction for each locus in the cluster. For optimization, we use a loss function that penalizes the squared difference between the observed allele fractions and the cluster allele fractions (methods). We solve this with the optimized gradient descent algorithm ADAM^13^ using tensorflow^14^. The advantage of mixture model clustering over hard clustering is that cells can be partially assigned to multiple clusters, which naturally allows for both doublet cells and varying levels of ambient RNA (Fig. 1d). Having obtained the cluster centers, we identify doublet cell barcodes (Fig. 1e) by modeling a cell’s allele counts as being drawn from a beta-binomial distribution whose parameters are derived from either one or two clusters.

To identify the diploid genotypes of each cluster and the amount of ambient RNA (Fig. 1f), we assume that the allele counts for locus *i* of each cluster *j* are drawn from a binomial distribution with an alternative allele fraction of (1-*ρ*)*f*_*ij*_+*ρ*\**a*_*i*_, where *f*_*ij*_ is 0, 0.5, or 1 (with a haploid mode limited to 0 and 1), *ρ* is a parameter representing the amount of ambient RNA and *a*_*i*_ is the average allele fraction in the experiment. The ambient RNA shifts the observed allele fraction away from the underlying genotype allele fractions^3^. This model is implemented in the domain-specific language for probabilistic models, STAN^15^, and it solves for the maximum likelihood soup fraction with gradient descent.

There has been some concern in the community that it will be difficult to know which cluster corresponds to which individual after deconvolution with multiplexed scRNAseq experiments when genotypes are not known a priori. To address this, we propose an experimental design involving *m* overlapping mixtures for *2*^*m*^*−1* multiplexed individuals (Table 1). Each individual is assigned a binary number from *1* to *2*^*m*^, where each bit corresponds to the inclusion (1) or exclusion (0) from each of the mixtures. This gives each individual a unique signature of inclusion/exclusion across the mixtures. Although each sample is in a different number of mixtures, the number of cells per experiment can be adjusted according to the number of mixtures that contain that sample.

**Table 1:** Sample-cluster deconvolution experimental design. This table outlines an experimental design of seven individuals with three overlapping mixtures to allow for clusters to be assigned to individuals. **a**, Shows the mapping of individuals to binary numbers where each digit of the binary number represents inclusion/exclusion from a mixture. **b**, The resulting mixtures.

**a. Binary mapping**
| Mixture | 1 | 2 | 3 |
| --- | --- | --- | --- |
| Individual a | 0 | 0 | 1 |
| Individual b | 0 | 1 | 0 |
| Individual c | 0 | 1 | 1 |
| Individual d | 1 | 0 | 0 |
| Individual e | 1 | 0 | 1 |
| Individual f | 1 | 1 | 0 |
| Individual g | 1 | 1 | 1 |

**b. Mixtures**
| Mixture 1 | d | e | f | g |
| --- | --- | --- | --- | --- |
| Mixture 2 | b | c | f | g |
| Mixture 3 | a | c | e | g |

Currently, there are no good generative models available for batch effects, allele-specific expression, ambient RNA, and doublets in scRNAseq that can be used to generate *in silico* data for testing methods that cluster by genotype. To generate realistic data with known ground truth we sequenced five lines of induced pluripotent stem cells (iPSCs) from the Human iPSC initiative^16^ with the 10x Chromium single cell system, both individually and in a mixture of all five lines (with three replicates of the mixture). Each mixture contained 5-7,000 cells and ~25,000 UMIs per cell (Table S1). We first synthetically mixed 20% of the cells from the 5 individual samples while retaining their sample of origin. To make the synthetic mixture as close to real data as possible, we also simulated 6% doublets by switching all of the reads’ barcodes from one cell to that of another cell and 5% ambient RNA by randomly switching cell barcodes for 5% of the reads. A low dimensional representation of the expression matrix, *E*, reveals relatively little variation as expected since there is only one cell type present (Fig. 2a). Indeed, the most significant driver of expression appears to be the donor of origin, but the donor cells overlap in expression patterns and it is not possible to assign a donor to each cell based solely on expression patterns.

**Figure 2:**
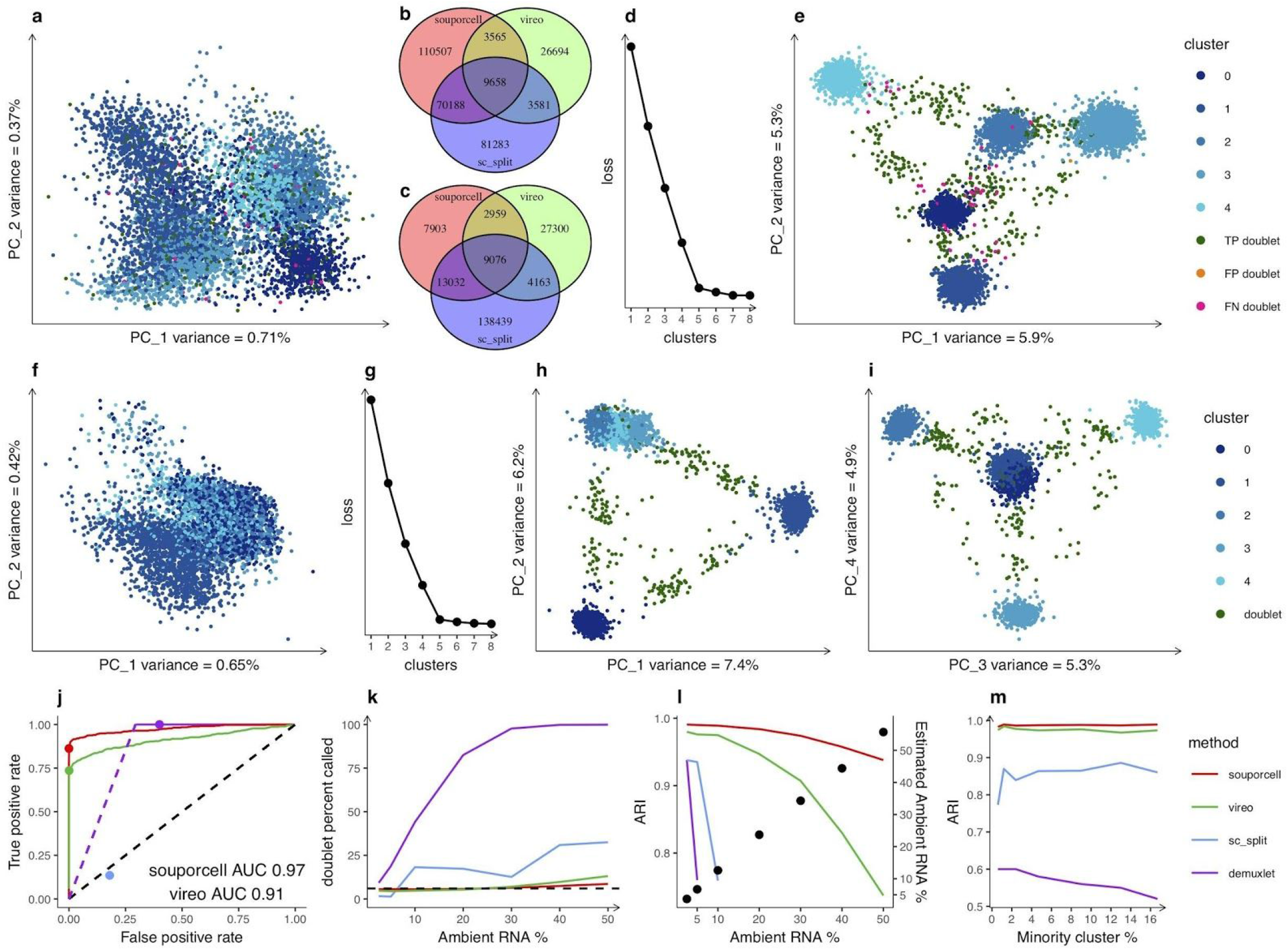
Evaluation of clustering accuracy. **a**, Expression PCA of a synthetic mixture of five HipSci cells lines with 5% ambient RNA and 6% doublets colored by known genotypes. Because these samples only contain one cell type, the largest remaining source of variation in the expression profile comes from the genotype, although the signal is not sufficient for accurate genotype clustering. **b**, Venn diagram of the number of total SNPs identified by each tool, and **c**, used by each tool for clustering. **d**, Elbow plot of the number of clusters versus the loss value showing a clear preference for the correct number of clusters (k=5). **e**, PCA of the normalized cell-by-cluster loss matrix from souporcell. As this is a synthetic mixture in which we know the ground truth, we color by genotype clusters and highlight errors. **f**, Expression PCA of a single replicate (see Fig. S1 for reps) of the experimental mixtures (4925 cells) colored by genotype clusters from souporcell. **g**, Elbow plot of the loss versus different numbers of clusters showing a clear preference for the correct number of clusters. **h** and **i**, PCAs showing the first four PCs of the normalized cell-by-cluster loss matrix colored by cluster. **j**, ROC curve of the doublet calls made by souporcell and vireo and a point estimate for sc_split (blue dot) for a synthetic mixture with 6% doublets 451/7073 and 10% ambient RNA. We show both the curves and the threshold chosen (points) for each tool. Sc_split did not give a score so we simply show the point estimate. Demuxlet’s doublet probabilities were all 1.0 until the solid line starts, so we show a theoretical dotted line up to that point. **k**, Doublet call percentages for all tools on synthetic mixtures for varying amounts of ambient RNA versus the actual doublet rate (dotted line). **l**, Adjusted Rand Index (ARI) versus the known ground truth of synthetic mixtures with 6% doublets and a varying amount of ambient RNA. For levels >=10% ambient RNA, sc_split identified one of the singleton clusters as the doublet cluster, which means that the ARI was not clearly interpretable. Right y-axis vs points shows the estimated ambient RNA percent by souporcell versus the simulated ambient RNA percent. **m**, ARI of each tool on a synthetic mixture with 8% ambient RNA and 6% doublet rate with 1,000 cells per cluster for the first four clusters and a variable number of cells in the minority cluster (25-800 cells in the minority cluster).

We compared souporcell to vireo and sc_split by first running variant calling and cell allele counting as recommended for each tool (methods). The number of SNPs reported and used for clustering (Fig. 2b, c) vary dramatically across the three methods. Using souporcell, we clustered cells by their genotypes, and evaluated the correct number of clusters through an elbow plot comparing the loss versus a varying number of clusters (Fig. 2d). The clustering output can be viewed as a matrix with cells as rows and clusters as columns with the values being the loss of that cell versus the corresponding cluster. To visualize the five clusters identified by genotype we carried out a Principal Component Analysis (PCA) of the normalized loss matrix, which reveals a clear separation of the clusters, with interspersed doublets (Fig. 2e). For these data souporcell assigned 7017/7073 singletons and 396/451 doublets correctly; one singleton was falsely labeled as a doublet, and 55 doublets were misidentified as singletons. We carried out the same analysis for the three replicates of the experiment mixtures and show results for one (Fig. 2f,g; see Fig. S1 for replicates). The expression PCA (Fig. 2f) and normalized cell-cluster loss PCA (Fig. h,i) of the experimental mixture were similar to the synthetic mixture indicating that the synthetic mixtures were an accurate approximation of real mixtures with the notable exception that while two PCs were sufficient to separate the genotypes in the synthetically mixed sample, the real mix requires four PCs. This discrepancy is most likely due to complexities of the real mixture that were not accounted for by our simulations. To compare doublet detection between methods, we calculated a receiver-operator characteristic (ROC) curve of the doublet calls (Fig. 2j) on a synthetic mixture with 6% doublets and 10% ambient RNA that showed the area under the curve values of 0.97 and 0.91 for souporcell and vireo, respectively. We also show point estimates for the doublet threshold chosen. Demuxlet’s posterior doublet probability output did not have enough significant digits and is 1.0 until it starts varying with 27% false positives. By default, a threshold with nearly 40% false positive doublets is chosen.

Each of the five human iPSC lines has existing WGS data generated as part of the HipSci Project^17^. Therefore, for the experimentally mixed replicates, we compared each tool’s clustering to sample assignments obtained from demuxlet using genotypes available from the WGS. Demuxlet significantly overestimates doublets versus expectations based on the number of cells loaded^9^ (Table S2) especially as ambient RNA increases (Fig. 2k). Because we could not trust the doublet calls of demuxlet, we allowed sc_split, vireo, and souporcell to exclude their called doublets and then compared the remaining cells to demuxlet’s best single genotype assignment. The Adjusted Rand Index (ARI) of the remaining cell assignments versus demuxlet (Table S2) were 1.0 (fully concordant) for souporcell and vireo across the three replicates and an average of 0.97 for sc_split.

To evaluate the robustness of each tool across a range of parameters, we created synthetic mixtures of the five individual human iPSC scRNAseq experiments to test both the sensitivity to the ambient RNA level (Fig. 2k, l) and the ability to accurately assign cells to a cluster if it is much smaller than other clusters (Fig. 2m). For the ambient RNA experiment, we synthetically combined 20% of the cells from each of the five individual samples and simulated 6% intergenotypic doublets and a range of ambient RNA from 2.5%-50% representing realistic ranges previously reported^3^. We found that souporcell and vireo retain high accuracy with souporcell being more robust at accurately calling doublets in high ambient RNA cases (Fig. 2l). The ARI of sc_split and demuxlet suffered due to poor doublet detection. With these data we also show that souporcell is able to accurately estimate the amount of ambient RNA in the experiment (Fig. 2l). To test robustness to sample skew, e.g., one donor’s cells are underrepresented, we created a set of synthetic mixtures with 1,000 cells from each of four individual samples and 25-800 cells for the minority cluster including 8% ambient RNA and 6% doublets (Fig. 2m). We found that all tools performed well down to the minority cell cluster comprising only 1.2% (50 cells) of total cells (Fig. 2m), but only souporcell and vireo were able to correctly identify all minority sample singletons as their own cluster down to 0.6% of all cells. Again, demuxlet’s poor ARI was due primarily to extremely high levels of false positive doublets (Fig. 2k).

We then compared souporcell’s genotype and ambient RNA co-inference to vireo and sc_split versus the variants called from whole genome sequencing data. As previously mentioned, the total variants available for genotyping differed between tools (Fig. 2b). In scRNAseq data most variants have very low coverage per cluster compared to what would be generated from WGS data, thus the genotype accuracy is significantly lower than one would attain with genome sequencing. Nevertheless, souporcell surpasses both vireo and sc_split in genotype accuracy on a synthetically mixed sample with 6% doublets and 10% ambient RNA (Fig. S1i). The most common error mode for vireo and sc_split is calling homozygous reference loci as heterozygous variants (Fig. S1j) which is expected when ambient RNA is not accounted for, as it is not in these two tools.

Next, we considered more challenging scenarios involving multiple cell types, widely varying numbers of cells per sample, and closely related genotypes. The decidua-placental interface plays an important role in pregnancy and birth, and is of importance to several diseases, including pre-eclampsia^19^. Recently, more than 70,000 cells were profiled by scRNAseq^18^ to explore the transcriptional landscape at this interface. The decidua is primarily composed of maternal cells with some invading fetal trophoblasts, while the placenta is largely composed of cells of fetal origin with the exception of maternal macrophages. In the study exploring this interface^18^, WGS from blood and placenta was used to genotype both mother and fetus, and demuxlet was used to assign cells to each individual. Here, we applied souporcell, vireo, and sc_split to two placental samples and one decidual sample from a single mother to determine if cellular origins could be established without reference genotypes. We show the expression t-SNE of a single placental sample labeled by cell type annotation^18^ and colored by genotype cluster as assigned by each method (Fig. 3a). While souporcell clusters agree with demuxlet and segregate with the expected cell type clusters, vireo and sc_split have major discordances with demuxlet. This is similar for the other samples tested (Fig. S2, Table S3). Comparing souporcell to demuxlet, there are 21 cells that demuxlet labels as maternal or fetal but which appear in the other individual’s cell type clusters. Based on the position of these cells in the expression t-SNE plot, it is most likely that these are errors in the demuxlet assignments that are not made by souporcell.

**Figure 3:**
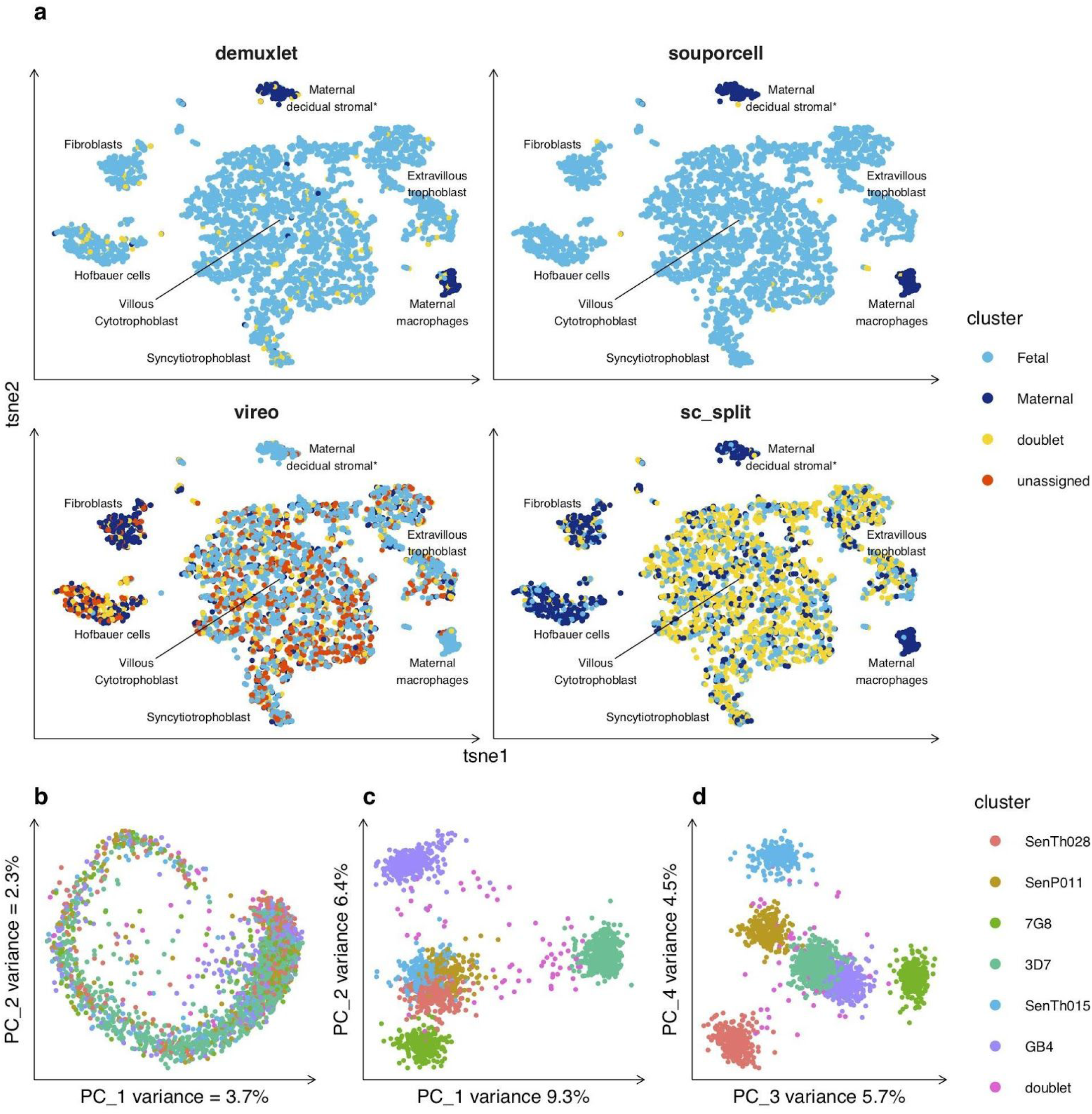
Application to challenging datasets. **a**, Cell expression t-SNE plots of 3,835 cells colored by each tool’s genotype assignments or clusters for placenta1 (other samples in Fig. S2). Cell phenotype clusters and cell genotype clusters co-segregate, with the majority of cell types being of fetal origin with the exception of maternal macrophages and *maternal decidual stromal cells, the latter of which (found only in one donor) were considered to be a non-placental artefact arising from the surgical procedure and were removed during data quality control in the original study^18^. We observe high concordance between souporcell and demuxlet (ARI 0.95) whereas vireo and sc_split have large discordances with ARI of 0 and 0.03 respectively. **b**, Expression PCA colored by genotype clusters for Plasmodium sample 1 (2608 cells) (other samples in Fig. S3) showing an even spread of genotypes throughout the asexual lifecycle. **c** and **d**, PCAs of first four PCs of souporcell’s normalized cell-by-cluster loss matrix showing good separation of each genotypic cluster.

We also tested souporcell on a non-human sample, the single-celled malaria parasite *Plasmodium falciparum*, for which single cell approaches are now used to explore natural infections^20^. Malaria infections often contain parasites from multiple different genetic backgrounds, and it is not possible to separate the strains prior to sequencing. These samples differ from human samples in a variety of ways; they are haploid when infecting humans, the genome is >80% A/T, and the transcriptome is only ~12 megabases (genome is ~23 Mb). We generated three datasets containing six genetically distinct strains of *P. falciparum* (methods) sampling 1893-2608 cells with median UMIs of ~1000 (1/25th of the HipSci samples). Analysis of the expression profile of one of these (see Fig. S3 for the others) reveals that the genotypes are evenly distributed across the *Plasmodium* intra-erythrocytic cycle (Fig. 3b) while being well separated in normalized loss cluster space (Fig. 3c,d). The ARI for each method (Table S4) on the three *Plasmodium* data sets show superior performance for souporcell across the board, with sc_split suffering on all datasets and vireo performing poorly on one, which had an ARI versus demuxlet of 0.24. This sample was more difficult due to sample skew caused by a clonal expansion of one of the six strains.

## Discussion

Here we have presented souporcell, a novel method for clustering scRNAseq cells by genotype using sparse mixture model clustering with explicit ambient RNA modeling. Our benchmarks show that souporcell can outperform all other currently available methods, including those that require genotypes *a priori*. Using more realistic and challenging test cases than previous studies, we show that souporcell is robust across a large range of parameters, and more so than any other currently available method. Moreover, souporcell is highly accurate for challenging datasets involving closely related maternal/fetal samples, and varying mixtures of *Plasmodium falciparum* strains. Due to the advantages that mixtures give to scRNAseq experiments in ameliorating batch effects, improving doublet detection, and allowing for ambient RNA estimation, souporcell enables donor multiplexing designs to be used more easily than was previously possible, including in situations when no WGS or genotyping data are available. In addition to reducing cost and allowing for more complex and robust experimental designs, souporcell also enables valuable genotype information to be extracted and ambient RNA estimation at no additional cost.

## Data Availability

Maternal Fetal data is available at https://www.ebi.ac.uk/arrayexpress/experiments/E-MTAB-6701/ with accession numbers FCA7474063-FCA7474065. This data is shown in Fig 3, Supp Fig 2. HipSci cell line data will be made available on GEO upon acceptance. This data is shown in Fig 2, Supp Fig 1. Plasmodium data will be made available on GEO upon acceptance. This data is shown in Fig 3, Supp Fig 3.

## Code Availability

Souporcell is freely available under the MIT open source license at https://github.com/wheaton5/souporcell.

## Acknowledgments

We acknowledge the Wellcome Sanger Institute’s DNA Pipelines for construction of the 10x sequencing libraries. We thank Allan Muhwezi and Andrew Russell for assistance with parasite culture and 10x Single-cell 3’ RNA-seq respectively. In addition, we would like to thank Matthew Young for useful conversations about ambient RNA, Mirjana Efremova for providing information about the maternal/fetal data, and Katie Gray for assistance in interpreting the previously unannotated cluster.

## Author contributions

MKNL and MH conceived of the project. HH developed the methods, software, ran the tests and simulations, and created the figures. MKNL, MH, and HH wrote the manuscript with methods contributions from AT, AK, and MI. AT conducted the *Plasmodium* wet lab experiments. AK and MI conducted the HipSci cell line experiments. RD provided feedback and guidance throughout the project.

## Supplementary Methods

### Remapping

We remap reads due to several different artifacts, described below. We first take the STAR aligned bam and create a fastq file from it using pysam and a custom python script (available at https://github.com/wheaton5/souporcell/renamer.py) while placing the UMI and cell barcode information in the read name for later use. We map these reads to the reference genome using minimap2 version 2.7-r654 with parameters -ax splice -t 8 -G50k -k 21 -w 11 --sr -A2 -B8 -O12,32 -E2,1 -r200 -p.5 -N20 -f1000,5000 -n2 -m20 -s40 -g2000 -2K50m --secondary=no, but have seen similar accuracy with the RNAseq aligner HiSat2^21^. We resupply the cell barcode tags and UMI tags to the bam using pysam and a custom python script (available at https://github.com/wheaton5/souporcell/retag.py) and sort and index the bam file with samtools. All steps are now encapsulated into a simple pipeline script and provided as a singularity container for easy installation.

We identified three different artifacts introduced by the STAR alignments resulting in false positive variants as well as reference bias that causes reads that do not support the reference allele to appear as though they do. The first artifact is due to the way STAR handles spliced reads when the read does not match the reference well. STAR will take such a read and introduce multiple splice events that force it to fit the reference in a statistically spurious fashion. We have observed cigar strings such as 8M129384N12M50238N77M where the read matches one location well with 77 matches plus mismatches in one location. Instead of soft clipping the initial 20 bases that did not have a statistically significant alignment, STAR introduced two large splicing events to match very short regions (8 and 12 bases) to the reference. Due to the limitation of a read having a single mapping quality and these spliced reads being encoded as a single read object in the bam file, the variant callers will treat these spurious matches as high mapping quality. Consequently, for variants in these loci, the variant callers will count these reads as supporting the reference allele, thereby introducing reference bias and noise to the downstream clustering. This has been noted by others in the past, and GATK recommendations for variant calling on bulk RNAseq https://gatkforums.broadinstitute.org/gatk/discussion/3891/calling-variants-in-rnaseq involve removing these regions of the alignments prior to variant calling. The second set of artifacts is alignment parameter differences between STAR and aligners intended for variant calling. The second type of artifact we found was due to the soft clip penalty being higher in STAR and not being exposed as a parameter to the user. This leads to false positive variants due to the lack of soft clipping where other mappers would soft clip poorly matching read ends. The final issue is that the indel penalty relative to the mismatch penalty is much higher in STAR than other aligners. This causes the alignments to choose many mismatches over a single or few indels when possible and thus create false positive variants. This is a parameter which is exposed to the user but, the default makes the output of cellranger poorly suited for variant calling. For these reasons, we find it best to remap these reads with a mapper specifically tuned to genomic variant calling.

### Variant Calling

#### Souporcell

Variant calling consists of two steps. First we identify candidate SNPs using freebayes (version v1.3.1-17-gaa2ace8) with parameters -iXu -C 2 -q 20 -n 3 -E 1 -m 30 --min-coverage 6 --max-coverage 100000 --pooled-continuous. If one wished to use known common variant sites, one could skip this step and provide that vcf to the following step. In the second step we count alleles for each cell using the program vartrix (available at https://github.com/10XGenomics/vartrix (release version 1.1.3)) with parameters --umi --mapq 30 --scoring-method coverage which gives us two sparse matrix outputs which represent the UMI allele counts per cell for each locus. For souporcell, we limit the loci considered for clustering to the ones with at least *n* cells (default =10) supporting each allele. For all human samples we used 10, but for the *Plasmodium* samples, we used *n*=4 due to the lower number of variants in the *Plasmodium* data. This provides us with fairly robust SNPs that have a good chance of aiding the clustering process.

#### Vireo

Vireo recommends running cellSNP (https://github.com/huangyh09/cellSNP version 0.1.6) on the STAR aligned bam with parameters --minMAF 0.1 --minCOUNT 100 limiting the analysis to loci with at least 100 UMIs and 10% minor allele fraction which are the settings we used throughout our analysis. For vireo donor clustering we use their R package, cardelino version 0.3.8.

#### Sc_split

Sc_split recommends using freebayes (the version we ran test on was v1.3.1-17-gaa2ace8) on the STAR aligned bam with parameters -iXu -C 2 -q 20 and then filtering for SNPs with a quality score >=30. We used bcftools for filtering with the command bcftools filter -e ‘QUAL<30’. Then this vcf is used along with the matrix.py script in sc_split with the filtered vcf, the STAR aligned bam, and the cell barcode file as input to get the allele counts for each cell. For sc_split donor clustering we used git commit hash 52face6a4c1b291651bdf9b56328d168c7cb1fa6 cloned from master at https://github.com/jon-xu/scSplit on April 21, 2019.

### Sparse mixture model clustering

Definitions

- *K*: number of genotype clusters to be fixed at the outset. Lower case *k* will be used for indexing and referring to a specific cluster.
- *C*: number of cells. Lower case *c* will be used for indexing and referring to a specific cell barcode. This barcode could have 0, 1, or more cells. It is important for some assumptions in this model that the majority of barcodes contain a single cell.
- *L*: number of variant loci. Lower case *l* will be used to index and refer to a specific locus. We will assume only biallelic variants. *L*_*c*_ will be a list of loci with observed data in cell *c*.
- *A*: Allele counts. *A*_*l,c*_ is a vector of size 2 with the first number representing the number of reference alleles and the second value representing the number of alt alleles seen at locus *l* in cell *c*.
- *ϕ*_*k*,*l*_: mixture parameter for allele fractions of cluster *k* at locus *l*. This is a real number representing the fraction of ref alleles in this cluster at this locus. We expect this to be near 1.0 (homozygous reference), 0.5 (heterozygous), or 0.0 (homozygous alt) but will be skewed from these values by noise, doublets, and ambient RNA.

#### Model

We define a loss function that will penalize the squared difference of the allele fractions of a cell and the cluster to which it is closest.

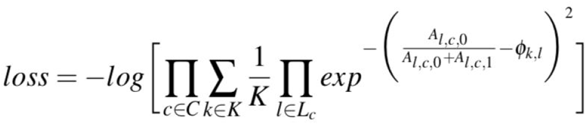

We minimize this loss function using tensorflow and the ADAM optimizer with a learning rate of 0.1 and a maximum epochs of 1000 with convergence criteria of a loss difference over 10 epochs of less than 1. We randomly initialize cluster centers and run this optimization 15 times by default to avoid local minima and take the solution with the minimum loss. While this loss function is not a log probability, we treat it as such and then define the pseudo-posterior for cell assignments to clusters as the following.

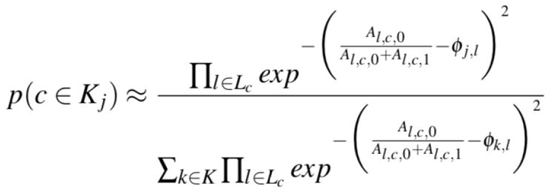

### Doublet detection

Definitions

- *A*_*k*,*l*_: Allele counts at locus *l* for all cells in cluster *k* according to the maximum probability cluster assignment from our clustering. This is a vector of size two with the ref and alt allele counts.

We treat the allele counts of each cell at each locus as random variables drawn from a beta-binomial distribution from either a single cluster or a pair of clusters. The beta-binomial is used to model our uncertainty in the binomial parameter *p*. For a single cluster the parameters are alpha = 1+alt counts and beta = 1+ref counts.

For the singleton case, we have

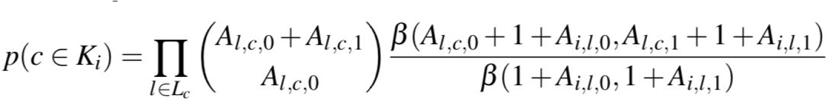

Where *β* is the beta function and cluster *i* is the best fitting cluster for cell *c*.

The expected allele fractions of a doublet coming from cluster *i*, and cluster *j* is the average of the allele fractions of the two clusters. To obtain the pseudocounts needed to parameterize the beta-binomial, we use the total counts of the cluster with less coverage at this locus. That is,

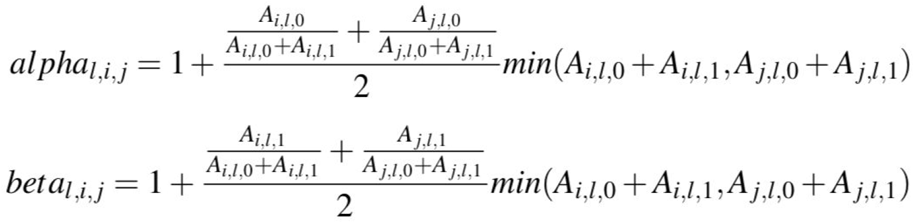

The doublet probability given those conservative parameters becomes

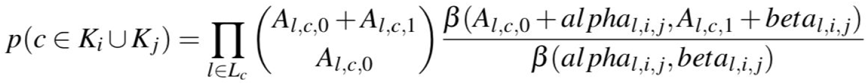

We only consider the possibility of a cell coming from its first best cluster as a singleton or its top two matching clusters as a doublet. This assumption provides both simplicity and speed, and we have found it to be adequate. The posterior for each cell being a doublet is then given by

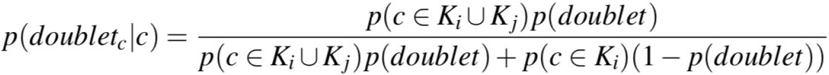

Where cluster *i* is the best fitting cluster for cell *c* and cluster *j* is the second best fitting cluster for cell *c*. We allow the prior to be set by the user but have used an uninformed prior of 0.5 for all of our analysis.

### Genotype and ambient RNA co-inference

Definitions

- *ρ*: mixture parameter representing the probability any given allele is arising from ambient RNA as opposed to from the cell associated with that barcode.
- *P*: ploidy. We assume ploidy is limited to 1 or 2.
- *A*_*l*_: total allele expression at locus *l*. This is again a vector of length 2 denoting the reference and alternative allele counts.
- *g*: used to denote the number of copies of the reference allele. The expected reference allele rate without ambient RNA is *g* and *g* `is an integer value ∈ [0`..*P*]. Note that for biallelic variants and ploidy 1 or 2, *g* is sufficient to uniquely determine the genotype.
- *p(true)*: prior for variant being a true variant vs a false positive. The default is 0.9 which was the value used for all analyses.

Here, the proportion of ambient RNA in the system, *ρ*, is the only free parameter and we solve for it using maximum likelihood. The model treats each locus in each cluster as coming from one of three genotypes for diploid (0/0, 0/1, 1/1, here denoted by *g* = 0, 1, or 2) and two genotypes from haploid (0, 1). We treat each cluster as independent and each locus as independent, before marginalizing across the possible genotypes. The model also considers the possibility of the variant being a false positive. In this case, the variant will not segregate into distinct allele frequencies between different clusters and it will most likely not attain a value close to the standard allele frequencies expected from the diploid or haploid genotypes. Thus, we model the allele counts in each cluster as having come from a mixture of ambient RNA (an average allele fraction in the experiment) and from the cells in that cluster. The observed allele fractions are assumed to have been drawn from a binomial distribution with a probability that was skewed away from *p* = *g/P* by the level of ambient RNA *ρ*. Thus, the probability of the binomial from which the allele counts are drawn for true positive variants is the following.

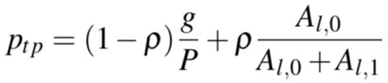

For a false positive the parameter is

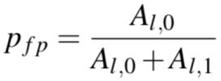

Thus, the full model is

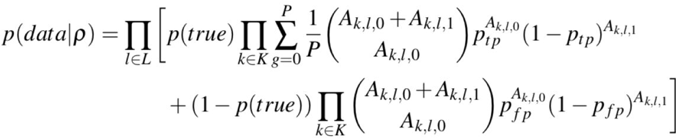

We solve for *ρ* with gradient descent using the statistical modeling domain specific language STAN. Next, we calculate the posterior of the variant being a true positive for each of the three (or two in the haploid case) genotypes versus it being a false positive. The prior on variants being true positives can be set by the user, but defaults to 0.9 which is the value used in our analyses.

### Human iPSC experiments

#### iPSC culture

Feeder-free iPSCs were obtained from the HipSci project^17^. Lines were thawed onto tissue culture-treated plates (Corning, 3516) coated with 5 μg/mL Vitronectin (rhVTN-N) (Gibco, A14700) using complete Essential 8 (E8) medium (StemCell Technologies, 05990) and 10 μM Rock inhibitor (Sigma, Y0503-1MG). Cells were propagated in E8 for 2 passages using 0.5 μM EDTA pH 8.0 (Invitrogen, 15575-038) for cell dissociation. Colonies were then dissociated into single cells using Accutase (Millipore, SCR005) and pooled in equal numbers, alongside individual lines, for one passage.

#### 10x Single-cell 3’ RNA-seq

To create a single cell suspension, iPS cells were cultured as described above in six-well plates before being washed once with room temperature D-PBS (Gibco, 14190-144). The D-PBS was removed before adding 1 mL of Accutase (Millipore, SCR005). The cells were incubated at 37°C for seven minutes before adding 1 mL of E8 media. The cells were collected in a 15 mL Falcon tube and triturated three times with a 5 mL stripette to obtain a single cell suspension. To ensure no cell clumps remained, the cell suspension was passed through a 40 μm cell strainer. The cells were counted and the viability was assessed on a Countess automated cell counter (Life Technologies). GEMs (gel beads in emulsion) were created using the 10x Genomics Chromium™ Controller, according to the manufacturer’s protocol. All channels were loaded such that an estimated 10,000 cells were captured for GEM formation and successful library preparation. All samples were processed using a 10x Genomics Chromium™ Single Cell 3’ v2 kit (PN-120237), following the manufacturer’s instructions. Libraries were multiplexed and sequenced at a rate of one library per lane of a Hiseq 4000 (Illumina), acquiring 150 bp paired-end reads.

**Supp Figure 1.**
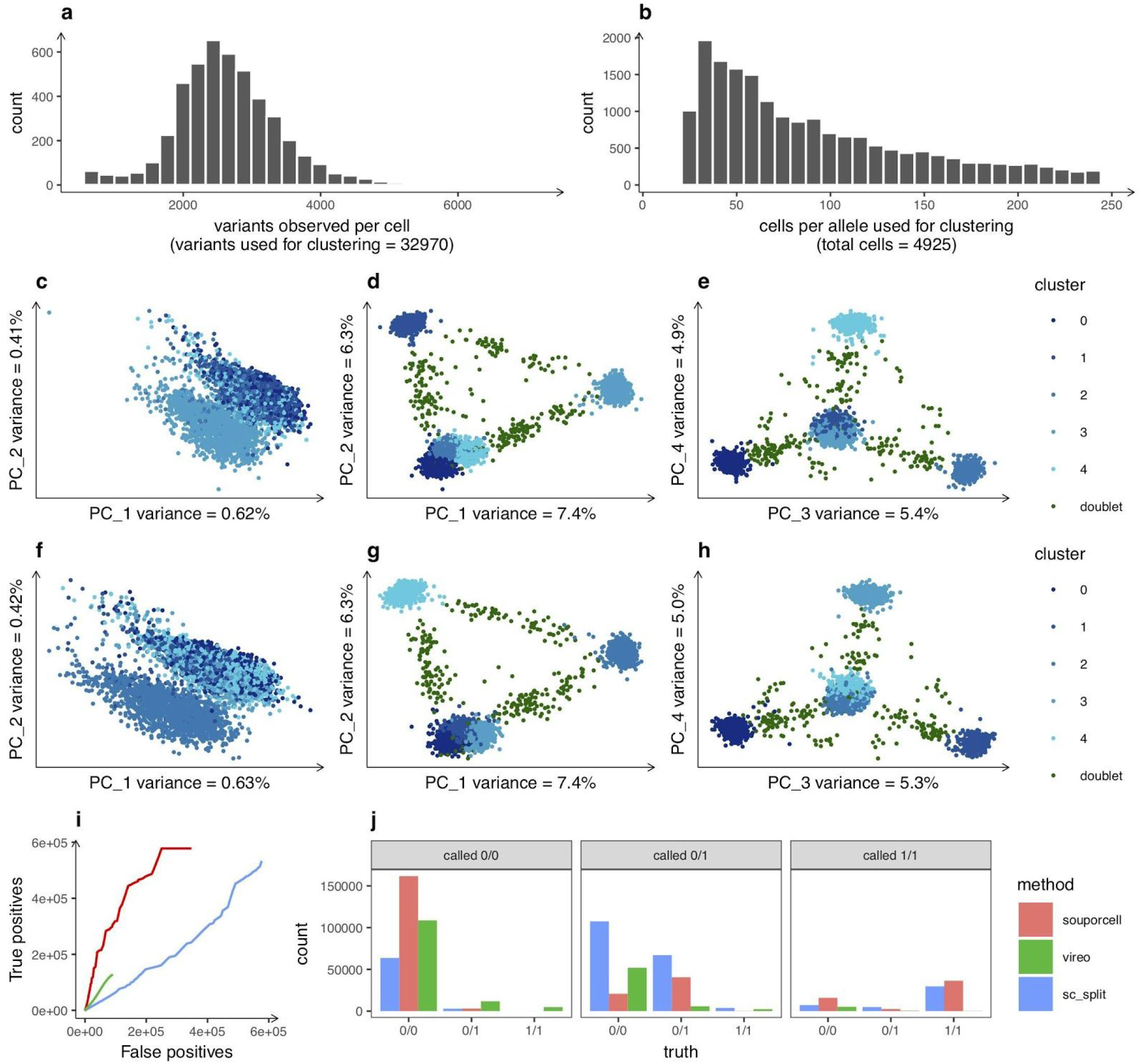
HipSci data sparsity, replicates, and genotyping. **a**, Distribution of the number of cells expressing a variant as well as **b**, the distribution of the number of alleles observed per cell that were used in souporcell clustering for HipSci mixture replicate 1 (replicates 2 and 3 are very similar, so not shown). **c**, Expression PCA of HipSci mixture replicate 2 (4832 cells) colored by genotype clusters from souporcell. **d**, and **e**, PCAs of the normalized cell-by-cluster loss matrix of HipSci mixture replicate 2 also colored by genotype cluster. **f**, Expression PCA of HipSci mixture replicate 3 (5144 cells) colored by genotype clusters. **g** and **h**, PCAs of normalized cell-by-cluster loss matrix of HipSci mixture replicate 3 colored by genotype cluster. **i**, Assessing genotype calling across souporcell, vireo, and sc_split. We plot true positive versus false positive genotype calls while sweeping the threshold on genotype likelihood. These are compared to a truth set obtained from variant calls on the WGS data **j**, Each method’s genotype calls versus the true genotype of each tool for a synthetic mixture of five HipSci lines with 6% doublets and 10% ambient RNA with a 0.95 probability threshold for each tool. The facets are the genotype calls made by each tool and the x-axis shows the correct assignments according to the WGS data. We observe that a major error mode for both vireo and sc_split compared to souporcell is that homozygous reference variants are mis-called as heterozygous because ambient RNA is not accounted for in these methods.

### Synthetic mixtures

We generated synthetic mixtures with custom python scripts using pysam and numpy. We took all of the reads for a subset of cell barcodes from each of the individual experiments and combined them into a new dataset. We then simulated doublet formation by randomly choosing among the cell barcodes that we had already chosen for the mixture experiment and then chose a cross-genotype cell barcode with which to create a doublet. We then took all of the reads of one of those cell barcodes and changed their cell barcodes to that of the other cell. We also simulated ambient RNA by randomly changing a read’s cell barcodes to that of another cell barcode at a specified rate. The values of each of these parameters are described in the text and figure captions.

### Demuxlet

We ran demuxlet git hash 85dca0a4d648d18e6b240a2298672394fe10c6e6 with default parameters except --field GT versus the cellranger bam, barcodes file, and vcf made by first downloading the exome bams from http://www.hipsci.org/ for each cell line, creating fastq files from them with samtools version 1.7 bam2fastq, then remapping to the cellranger reference with minimap2 version 2.7-r654 with parameters -ax sr, removing duplicates with samtools rmdup, and calling variants across the five bams with freebayes version v1.2.0-2-g29c4002-dirty with default parameters. Variants were then filtered with a custom python script using pyvcf such that the remaining variants be SNPs with QUAL >= 30.

### Maternal/Fetal

We obtained two placental samples and one decidual sample from Vento et al^18^ at https://www.ebi.ac.uk/arrayexpress/experiments/E-MTAB-6701/samples/. The samples used were FCA7474065 (placenta1) (Fig. 3a), FCA747064 (placenta2 (Fig. S2b)), and FCA747063 (decidua1 (Fig. S2a) all from the same individual. We obtained the fastq files and ran cellranger version 2.1.1 on them with default parameters to obtain the bam and cell barcodes files which are the input to our system. We then ran souporcell, sc_split, and vireo on them with recommended settings for each tool as previously detailed and obtained the demuxlet calls used in Vento et al^18^. We ran souporcell, vireo, and sc_split on each of these and compared them to the demuxlet calls excluding the demuxlet doublet cells and the doublets called by each tool (Supp Table 3).

**Supp Figure 2.**
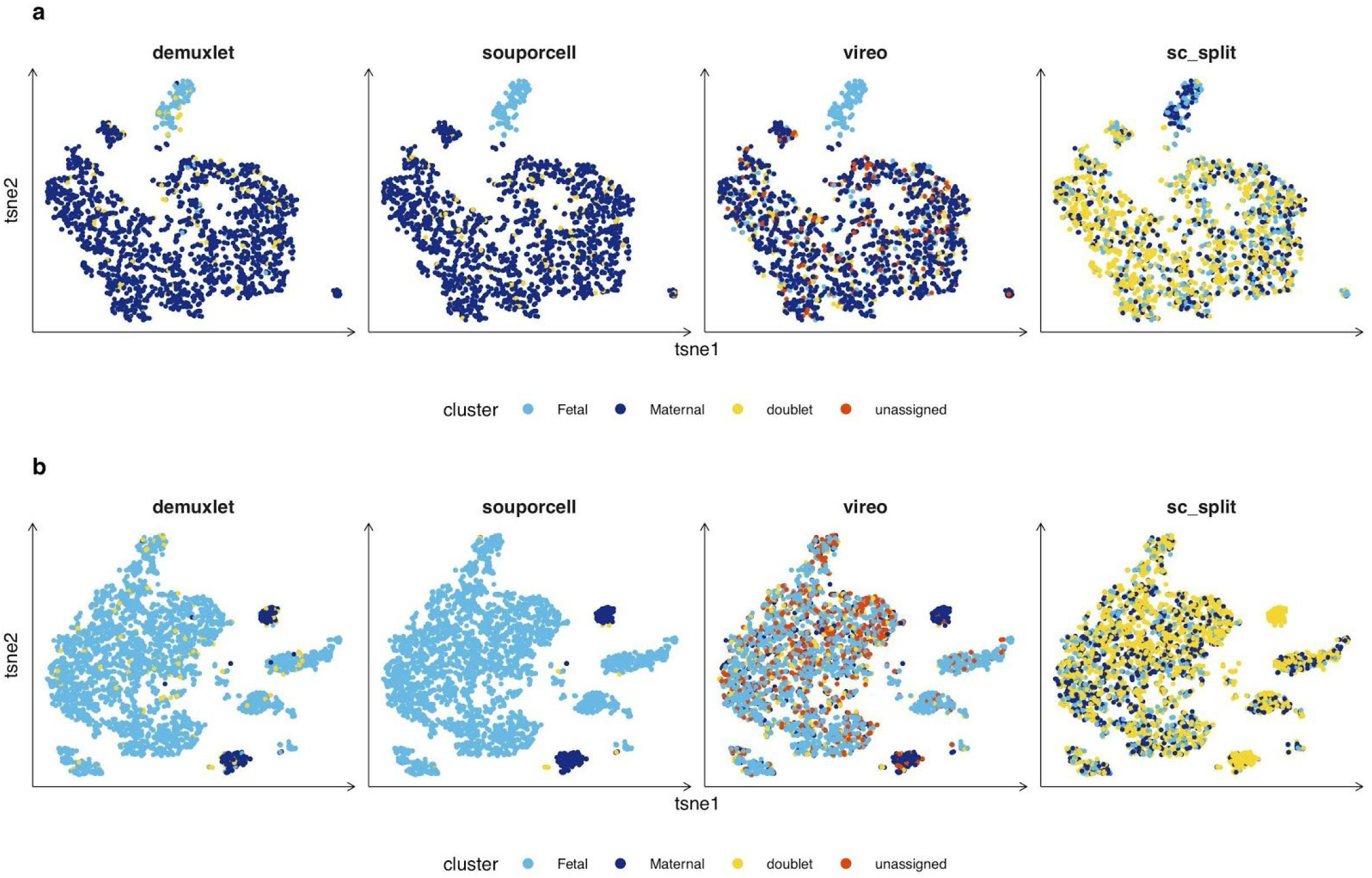
Maternal/Fetal decidua1 and placenta2. **a**, Expression t-SNE of a decidua1 sample (FCA747063, 2119 cells) colored by genotype clusters for each tool. Souporcell and demuxlet are highly concordant (ARI = 0.93). Vireo misidentifies a significant number of maternal cells as fetal cells. Excluding doublets and unassigned cells, vireo has an ARI of 0.3 versus demuxlet. Sc_split has many errors resulting in an ARI versus demuxlet of 0. **b**, Expression t-SNE of placenta2 sample (3968 cells) colored by genotype clusters for each tool. Souporcell is again highly concordant with demuxlet (ARI = 0.96). Vireo has significant problems producing an ARI vs demuxlet of 0.18, even when excluding doublets and unassigned cells called by either tool. Like the other maternal/fetal samples, sc_split struggles and has and ARI versus demuxlet of 0.

### *Plasmodium falciparum in vitro* culturing and single cell analysis

*P. falciparum* strains were maintained in O+ blood in RPMI 1640 culture medium (GIBCO) supplemented with 25 mM HEPES (SIGMA), 10 mM D-Glucose (SIGMA), 50 mg/L hypoxanthine (SIGMA), and 10% human serum in a gas mix containing 5% O_2_, 5% CO_2_ and 90% N_2_. Human O+ erythrocytes were obtained from NHS Blood and Transplant, Cambridge, UK. All samples were anonymous. *Plasmodium* culture using erythrocytes and serum from human donors was approved by the NHS Cambridgeshire 4 Research Ethics Committee (REC reference 15/EE/0253) and the Wellcome Sanger Institute Human Materials and Data Management Committee.

All *P. falciparum* clonal strains were obtained from MR4 (BEI resources): 3D7-HT-GFP (MRA-1029), 7G8 (MRA-152), GB4 (MRA-925), SenP011.02 (MRA-1176), SenTh015.04 (MRA-1181) and SenTh028.04 (MRA-1184). All strains were maintained in culture below 5% parasitemia for no less than 6 weeks without synchronization prior to the experiment in order to ensure maximum asynchronicity. Plasmodium1 pool was composed of 2 independently cultured flasks for each of the 6 strains. The Plasmodium1 pool was washed once in PBS, before resuspension in PBS at a concentration of 11,200 RBC/μl (corresponding to 479 parasites/μl). The Plasmodium2 pool was derived from an aliquot of the Plasmodium1 sample that had been resuspended in 200 μl of PBS and fixed with 800 μl of ice-cold methanol for 10 minutes on ice, before being washed twice in PBS and resuspended at 12,200 RBC/μl (corresponding 522 parasites/μl). The Plasmodium3 sample was derived from a mix of the 6 strains, grown in the same flask for 7 days and resuspended at 19,800 RBC/μl (corresponding to 960 parasites/μl). Hematocrits were established with a hemocytometer. Each cell suspension was loaded on one inlet of a 10x chromium chip according to manufacturer’s instructions with a target recovery of 9000 cells per inlet. Chromium 10x v2 chemistry was used and libraries were prepared according to the manufacturer’s instructions. Each 10x input library was sequenced on both lanes of a Hiseq 2500 Rapid Run using 75 bp paired-end sequencing.

**Supp Figure 3.**
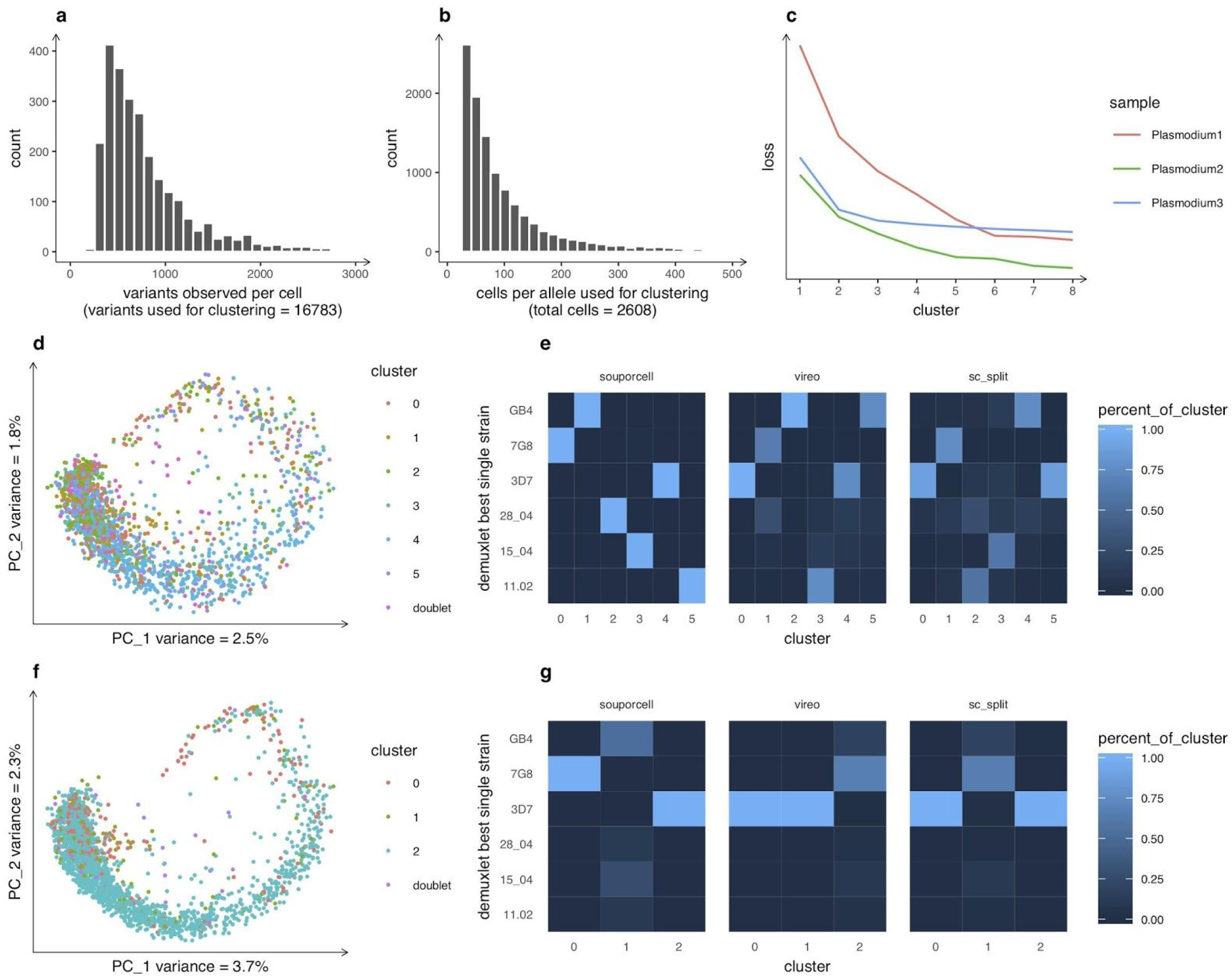
Plasmodium clustering. **a**, Distribution of number of variants observed per cell used for clustering (with at least 4 cells required to support each allele) and the total number of variants used for clustering on the Plasmodium1 sample. **b**, Distribution of counts of the number of cells expressing each allele used for clustering as well as the total number of cells in the Plasmodium1 sample. **c**, Elbow plots for each Plasmodium data set show relatively strong support for the correct number of clusters (6) for Plasmodium1, but less clear results for Plasmodium2, which suffered from higher amounts of ambient RNA, and for Plasmodium3, which due to more cell numbers biased towards three genotypes rather than a relatively even mixture. For this reason, we analyze Plasmodium3 with k=3. **d**, Expression PCA of the Plasmodium2 sample (1893 cells) colored by genotype clusters as called by souporcell. **e**, Confusion matrix heatmap of the demuxlet best single strain (Y axis) versus souporcell, vireo, and sc_split. For souporcell we see one cluster per strain as expected. Both vireo and sc_split have the majority strain, 3D7, split across two clusters and two other strains combined into a single cluster. **f**, Expression PCA of the Plasmodium3 sample (2293 cells) colored by genotype clusters as called by souporcell. **g**, Confusion matrix heatmap of the demuxlet best single strain (Y axis) versus souporcell, vireo, and sc_split genotype clusters with k=3. Souporcell clusters out the 3D7 and 7G8 strains correctly and puts all other cells into the final cluster while both vireo and sc_split put 3D7 into two clusters and all other cells into the remaining cluster.

### Software versions

Initial data analysis was done by cellranger v2.1.1 as input to souporcell, demuxlet, vireo, and sc_split

#### Souporcell

https://github.com/wheaton5/souporcell

We provide a singularity container build encapsulating all requirements for souporcell as well as a singularity definition file to recreate this container. The following are the software versions for all software excluding software required to build the system.

Freebayes – v1.3.1-17-gaa2ace8

Pyvcf – 0.6.7

Pysam – 0.15.3

Numpy – 1.17.0

Scipy – 1.3.0

Tensorflow – 1.14.0

Pystan – 2.17.1.0

Pyfasta – 0.5.2

Htslib – 1.9

Samtools – 1.9

Bcftools – 1.9

Vartrix – 1.1.3

Minimap2 – 2.7-r654

Bedtools – 2.28.0

#### Demuxlet

https://github.com/statgen/demuxlet

git hash 85dca0a4d648d18e6b240a2298672394fe10c6e6

#### Vireo

Cardelino R package version 0.3.8

cellSNP (https://github.com/huangyh09/cellSNP version 0.1.6)

#### Sc_split

Freebayes – v1.3.1-17-gaa2ace8

https://github.com/jon-xu/scSplit git commit hash 52face6a4c1b291651bdf9b56328d168c7cb1fa6

**Supp Table 1:** Sample metrics.

| Sample | Cells | Median UMI/cell | NOTES |
| --- | --- | --- | --- |
| Mixture1 | 4925 | 25155 | HipSci |
| Mixture2 | 4832 | 24913 | HipSci |
| Mixture3 | 5144 | 24807 | HipSci |
| euts | 4859 | 25088 | HipSci |
| nufh | 6781 | 17254 | HipSci |
| babz | 12299 | 11343 | HipSci |
| oaqd | 7107 | 18315 | HipSci |
| ieki | 6586 | 21969 | HipSci |
| SyntheticMixture | 7073 | 18793 | synthetic HipSci mixture (5% ambient RNA, 6% doublets) |
| Placenta1 | 3835 | 18415 | Maternal Fetal |
| Placenta2 | 3968 | 18046 | Maternal Fetal |
| Decidua1 | 2119 | 23075 | Maternal Fetal |
| Plasmodium1 | 2608 | 995 | *Plasmodium falciparum* |
| Plasmodium2 | 1893 | 762 | *Plasmodium falciparum* |
| Plasmodium3 | 2293 | 1126 | *Plasmodium falciparum* |

Number of cells and median UMI/cell for each sample including one representative synthetic mixture.

**Supp Table 2:** Hipsci clustering.

|  | method | mixture1 | mixture2 | mixture3 | synthetic mixture |
| --- | --- | --- | --- | --- | --- |
| ARI vs demuxlet single best (excluding doublets called by each tool) | souporcell | 1 | 1 | 1 | n/a |
|  | vireo | 1 | 1 | 1 | n/a |
|  | sc_split | 0.97 | 0.97 | 0.98 | n/a |
| ARI vs truth (including doublets) | souporcell | n/a | n/a | n/a | 0.99 |
|  | demuxlet | n/a | n/a | n/a | 0.76 |
|  | vireo | n/a | n/a | n/a | 0.98 |
|  | sc_split | n/a | n/a | n/a | 0.94 |
| ambient RNA | souporcell | 2.70% | 3% | 2.70% | 6.70% |
|  | truth | n/a | n/a | n/a | 5% |
| doublets | souporcell | 302 (6.1%) | 304 (6.3%) | 307 (5.9%) | 397 (5.6%) |
|  | demuxlet | 1537 (31) | 1811 (37.5%) | 1596 (31.2%) | 1329 (18.7%) |
|  | vireo | 306 (6.2%) | 287 (5.9%) | 285 (5.5%) | 311 (4.4%) |
|  | sc_split | 127 (2.6%) | 107 (2.2) | 102 (2%) | 89 (1.2%) |
|  | truth | n/a | n/a | n/a | 451 (6.4%) |
| euts | souporcell | 1040 | 1071 | 1180 | 844 |
|  | demuxlet | 790 | 743 | 922 | 799 |
|  | vireo | 1039 | 1074 | 1185 | 876 |
|  | sc_split | 1076 | 1102 | 1211 | 856 |
|  | truth | n/a | n/a | n/a | 817 |
| nufh | souporcell | 660 | 663 | 692 | 1191 |
|  | demuxlet | 317 | 264 | 330 | 1037 |
|  | vireo | 660 | 665 | 694 | 1211 |
|  | sc_split | 707 | 721 | 766 | 1271 |
|  | truth | n/a | n/a | n/a | 1183 |
| babz | souporcell | 747 | 729 | 734 | 2241 |
|  | demuxlet | 522 | 459 | 504 | 1591 |
|  | vireo | 742 | 734 | 740 | 2241 |
|  | sc_split | 818 | 742 | 806 | 2263 |
|  | truth | n/a | n/a | n/a | 2242 |
| oaqd | souporcell | 1590 | 1475 | 1602 | 1255 |
|  | demuxlet | 1341 | 1172 | 1345 | 1227 |
|  | vireo | 1590 | 1481 | 1609 | 1261 |
|  | sc_split | 1587 | 1501 | 1600 | 1353 |
|  | truth | n/a | n/a | n/a | 1247 |
| **ieki** | souporcell | 586 | 590 | 629 | 1145 |
|  | demuxlet | 418 | 383 | 447 | 1090 |
|  | vireo | 586 | 590 | 630 | 1160 |
|  | sc_split | 610 | 659 | 659 | 1241 |
|  | truth | n/a | n/a | n/a | 1133 |
| **unassigned** | souporcell | 0 | 0 | 0 | 0 |
|  | demuxlet | 0 | 0 | 0 | 0 |
|  | vireo | 2 | 1 | 1 | 13 |
|  | sc_split | 0 | 0 | 0 | 0 |

Evaluation of clustering metrics for each tool on HipSci experimental mixtures as well as one representative synthetic mixture with 5% ambient RNA and 6% doublets.

**Supp Table 3:**
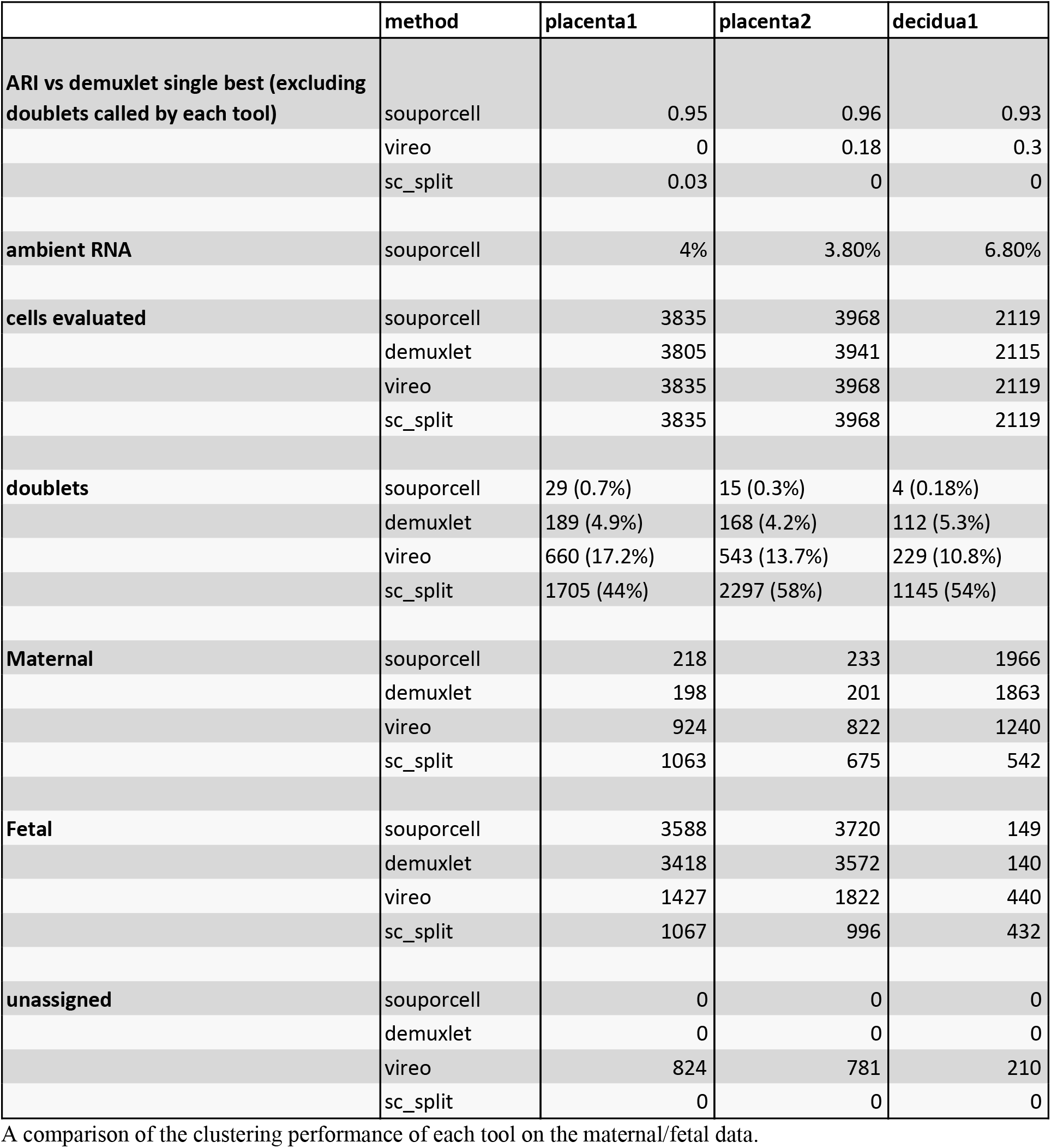
Maternal/Fetal clustering.

|  | method | placenta1 | placenta2 | decidua1 |
| --- | --- | --- | --- | --- |
| **ARI vs demuxlet single best (excluding doublets called by each tool)** | souporcell | 0.95 | 0.96 | 0.93 |
|  | vireo | 0 | 0.18 | 0.3 |
|  | sc_split | 0.03 | 0 | 0 |
| **ambient RNA** | souporcell | 4% | 3.80% | 6.80% |
| **cells evaluated** | souporcell | 3835 | 3968 | 2119 |
|  | demuxlet | 3805 | 3941 | 2115 |
|  | vireo | 3835 | 3968 | 2119 |
|  | sc_split | 3835 | 3968 | 2119 |
| **doublets** | souporcell | 29 (0.7%) | 15 (0.3%) | 4 (0.18%) |
|  | demuxlet | 189 (4.9%) | 168 (4.2%) | 112 (5.3%) |
|  | vireo | 660 (17.2%) | 543 (13.7%) | 229 (10.8%) |
|  | sc_split | 1705 (44%) | 2297 (58%) | 1145 (54%) |
| **Maternal** | souporcell | 218 | 233 | 1966 |
|  | demuxlet | 198 | 201 | 1863 |
|  | vireo | 924 | 822 | 1240 |
|  | sc_split | 1063 | 675 | 542 |
| **Fetal** | souporcell | 3588 | 3720 | 149 |
|  | demuxlet | 3418 | 3572 | 140 |
|  | vireo | 1427 | 1822 | 440 |
|  | sc_split | 1067 | 996 | 432 |
| **unassigned** | souporcell | 0 | 0 | 0 |
|  | demuxlet | 0 | 0 | 0 |
|  | vireo | 824 | 781 | 210 |
|  | sc_split | 0 | 0 | 0 |

A comparison of the clustering performance of each tool on the maternal/fetal data.

**Supp Table 4:**
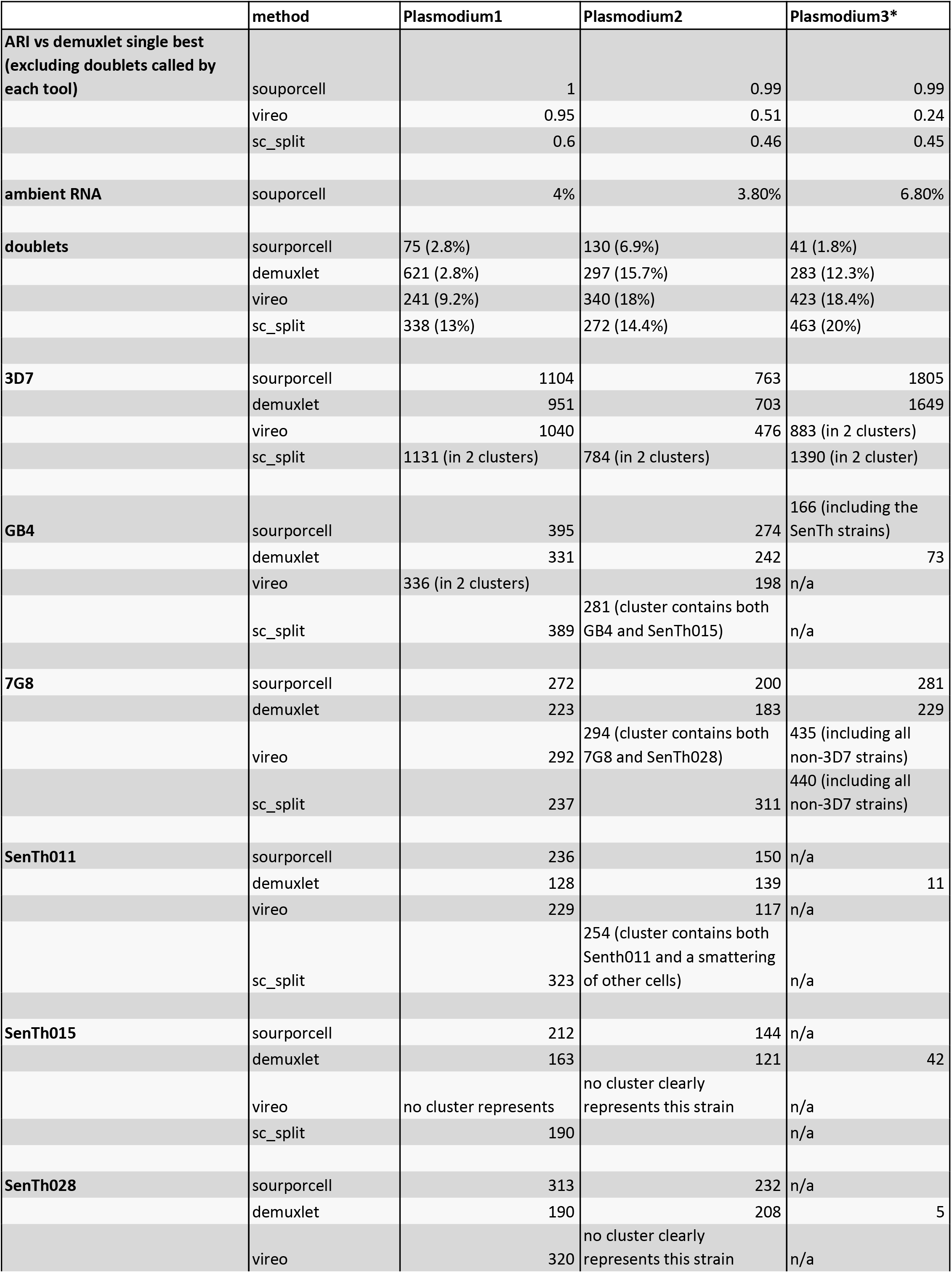
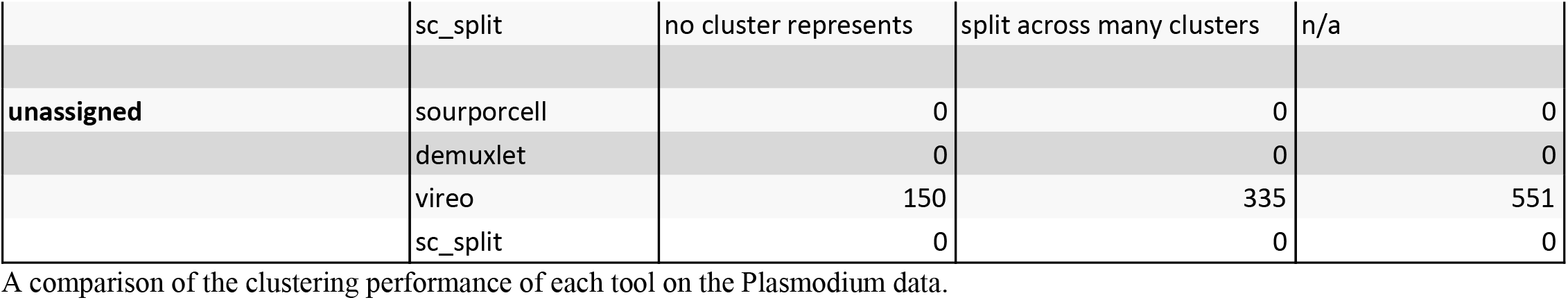
Plasmodium clustering.

